# Why resource dynamics matter in the optimization of microbial communities

**DOI:** 10.1101/2022.04.19.488725

**Authors:** Andrew D. Letten, William B. Ludington

**Affiliations:** School of Biological Sciences, University of Queensland, Brisbane, Queensland 4072, Australia; Department of Embryology, Carnegie Institution of Washington, Baltimore, MD, United States; Department of Biology, Johns Hopkins University, Baltimore, MD, United States

## Abstract

The impact of resource supply on microbial community dynamics is rarely treated explicitly in research into the control and optimization of natural and synthetic microbial communities. Using simulations, we show that compositional overlap between microbial communities decays rapidly with a change in the frequency of resource pulsing, particularly in the presence of a metabolic trade-off in resource affinity and maximum growth rate. We conclude that resource supply dynamics should be considered both a constraint and a tuning mechanism in microbial community optimization.

## Introduction

Understanding how biological communities assemble and the processes that stabilize them has been a central focus of ecology for over a century^1^. Recent decades have witnessed an accelerating impetus to leverage this fundamental understanding towards applied problems, from preserving biodiversity under anthropogenic environmental change^2^ to maintaining a healthy gut microbiota^3^. Microbes contribute to diverse physiological, biogeochemical and agricultural processes, and their experimental tractability holds promise for diverse interventions ranging from industrial and environmental remediation to human medicine and biofuel production^4,5^. However, microbial ecosystems have some fundamental differences from macro-scale ecosystems, and the conclusions drawn depend in large part on the way experiments and theory are conducted. Our thesis is that resource supply dynamics are fundamental to the control and optimisation of microbial ecosystems in nature and in the lab.

The modern study of microbiomes has been driven by advances in sequencing technology enabling researchers to describe microbial communities in extraordinary depth and to infer their operation through metabolic interactions. Building on this progress, the emerging field of synthetic microbial ecology seeks to develop a predictive theory of community assembly using mathematical and computational models integrated with experimentally-defined microbial communities in the lab. Theoretical approaches include those leveraging mathematical models and metabolic networks to predict which species combinations are stable and how they can optimize a given function (e.g., maximum biomass, waste degradation or host health)^6–10^. Experimental studies often take a combinatorial approach, iteratively assembling different species combinations *in vitro* and evaluating their stability and functional attributes ^11–14^. Both theory and experiments are valuable but they are also susceptible to their own *modus operandi* that may limit their correspondence and their translation to real world systems. On the one hand, theoretical approaches typically adopt the analytical tractability of steady state dynamics, where microbial consumers and the resources on which they depend are assumed to establish a stable equilibrium. On the other hand, experimental approaches almost exclusively embrace the high-throughput efficiency of serial-batch culture, where both the consumers and the resources on which they depend are made to fluctuate over several orders of magnitude with each serial passage. This raises two important questions: i) Should we expect unity in the composition of optimized communities deriving from continuous steady-state conditions (e.g., chemostat) vs the discontinuous dynamics of, for example, serial-batch culture? ii) How likely are optimized communities to persist under natural conditions? Here we argue that the answer to the first question is *rarely*, and that the answer to the second question will be highly contingent on how closely the resource dynamics adopted in theoretical and experimental work corresponded to those in nature.

To illustrate how different approaches to microbial community optimization can produce conflicting results, we focused on the difference between continuous versus pulsed resource supply. We performed simulations using a consumer-resource model, which tracks the abundances of species (consumers) and the resources they depend on through time. The growth rate of each species depends upon the availability of resources and the parameters of the species’ growth function. Bacteria often specialize on a specific resource, e.g. a carbon source, but can utilize other sources albeit with lower efficiency. Consumer-resource models account for these *substitutable* resources, and competition can lead to coexistence through specialization on alternative resources^15^. A key conclusion from such models is that under *steady-state* resource supply, the best competitors are those that can survive on the lowest concentration of one or more shared limiting resources. We set up our model to explicitly compare steady-state versus pulsed resource supply.

To set up the simulations, we randomly sampled the parameters of the Monod growth function, (*µ*_*max*_ [maximum growth rate] and *K*_*s*_ [half-saturation constant]) for five species competing for five substitutable resources. In one set of simulations (*n* = 100), we used both random *µ*_*max*_ and *K*_*s*_, and in another set we imposed a trade-off in maximum growth rate and substrate affinity 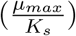 by making *K*_*s*_ = *µ*_*max*_^*4*^. The rationale for imposing a trade-off is that metabolic theory predicts that organisms that invest energy into a high maximum growth rate will have lower substrate affinities and vice versa ^16^. For each of the random competitor combinations, we simulated resources under continuous or pulsed resource supply with resource replenishment every 1/2, 1, 2, 4, 12 or 24 hours. The total resource flux was held constant under all frequencies of resource supply i.e., less frequent replenishment corresponds to larger resource pulses. Mortality was also held constant. After allowing the competitors to reach a steady state, which was time-averaged under pulsed treatments, we quantified the correspondence between the continuous supply treatment and the pulsed treatments using the Jaccard similarity index, 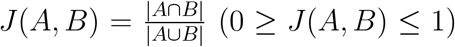, where the numerator gives the set of species (max = 5) that persist under continuous (*A*) *and* pulsed (*B*) resource supply, and the denominator gives the set of species (*max* = 5) that persist under continuous *or* pulsed resource supply.

A summary of the simulation results is provided in Figure 1. Under both sets of simulations (with and without imposing a rate-affinity trade-off) the similarity in final community composition between the continuous resource supply and the pulsed resource treatments decays with increasingly large intervals between resource replenishment. When no trade-off is imposed between *µ*_*max*_ and substrate affinity the mean compositional similarity is only 0.77 when resources are pulsed every 2 hours and down to 0.59 when resources are pulsed every 24 hours (typical of serial-batch culture). The rate of decay in the Jaccard index is more severe when a trade-off is imposed between maximum growth rate and substrate affinity, to the extent that once pulsing intervals reach four hours there is almost zero overlap in community composition. Ecological theory provides an intuitive explanation for the observed compositional dissimilarity between communities assembled under continuous and pulsed resource supply^17,18^. Examining the Monod function, 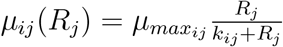, where 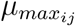 is the maximum growth rate and *k*_*ij*_ is the half saturation constant for consumer *i* on resource *j*, when resources are more continuously supplied, the better competitor is the one that can sustain a positive growth rate at the lowest concentrations of a limiting resource (i.e., has a higher substrate affinity or lower *R*^*\**^ in the language of resource competition theory^17^), which is greatly influenced by the *k*_*ij*_ term. In contrast, under increasingly pulsed resource supply, the better competitor is the one that can grow rapidly at higher resource concentrations. Having a low *R*^*\**^ is of little benefit if resource concentrations fluctuate over large amplitudes because it only confers an ephemeral competitive advantage in the brief period before the resource is completely depleted (ahead of the next resource pulse). Instead, a high maximum growth rate is optimal because it allows the consumer to grow rapidly and quickly deplete a shared limiting resource. This high maximum growth strategy is, however, sub-optimal under continuous resource supply because a low *R*^*\**^ strategist can draw the resource down and hold it at a concentration at which the maximum growth strategist is unable to maintain a positive growth rate. In our simulations, imposing this metabolically expected trade-off leads to a rapid decline in compositional similarity under increasingly large pulse intervals (blue line in Fig 1). We also observe a decline in compositional similarity when maximum growth rate and the Monod half saturation constant are randomly sampled independently of each other simply because the trade-off will emerge occasionally by chance (orange line in Fig 1). Two experimental tests of microbial community composition under continuous versus pulsed resource supply are consistent with these observations^19,20^.

**Figure 1:**
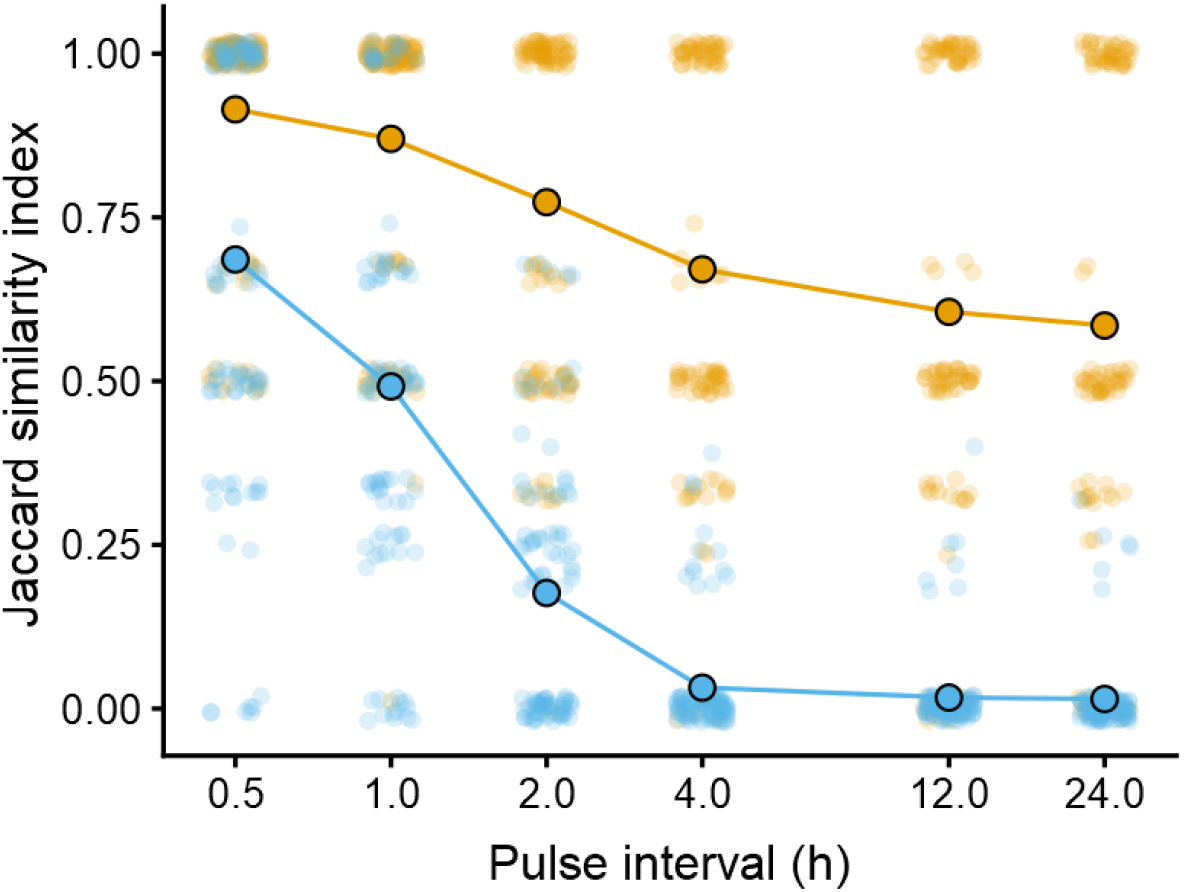
Compositional overlap (Jaccard similarity) between communities under continuous versus pulsed resource supply. Orange lines and circles denote model parametrisations with random sampling of both *µ*_*max*_ and *Ks*; blue lines and circles denote model parametrisations with *Ks* = *µ*_*max*_^*4*^. Small circles gives the result of an individual simulation; large circles indicate the corresponding mean.

The implications for optimization of microbial communities is that the resource supply regime needs to be tailored to the community being optimized. For example, wastewater treatment might be more appropriately modeled under continuous resource supply^21^, whereas fermented food and beverage production may be more closely allied to the pulsed resource dynamics observed in batch culture^22^. Resource supply might also be manipulated to favourably modify the competitive hierarchy in an existing community (e.g., by regulating the rate of nutrient supply to the gut). Indeed, there is emerging evidence that feeding frequency can drive significant changes in gut microbiota composition^23–25^. Thus, resource supply dynamics should be considered both a constraint in the design of novel microbial communities, and as a tuning mechanism for the optimization of pre-existing communities like those found in our own gut.

## Online methods

Simulation models take the general form:

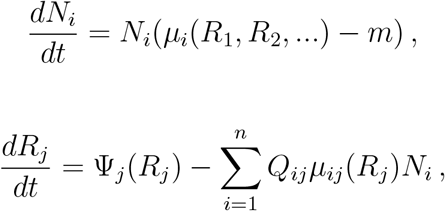

where *N*_*i*_ is the population density of consumer *i* (n = 5), *R*_*j*_ is the concentration of resource *j* (n = 5), *µ*_*i*_() is the per capita consumer functional response of consumer *i, d* is the per capita mortality rate / dilution rate, Ψ_*j*_(*R*_*j*_) is the resource supply function, and *Q*_*ij*_ is the resource quota of consumer *i* on resource *j* (amount of resource per unit consumer).

The consumer functional response is given by the Monod function, 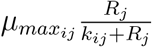, where 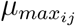 is the maximum growth rate and *k*_*ij*_ is the half saturation constant for consumer *i* on resource *j*.

Under chemostat dynamics, the resource supply function is given by Ψ_*j*_(*R*_*j*_) = *m*(*S*_*j*_ − *R*_*j*_), where *m* is the dilution rate and *S*_*j*_ is the supply concentration of resource *j*. Under pulsed resource dynamics Ψ_*j*_(*R*_*j*_) is replaced by discontinuous resource pulsing at fixed intervals: Δ*R*_*j*_ = *R*_*j*_ + *P*_*j*_, *t* = *kτ, k* = 1, 2, …,, where P is the size of the resource pulses, and *τ* is the pulse period (i.e., 0.5, 2, 4, 12, or 24 hours). In order to maintain an equivalent total resource flux across all treatments (continuous and pulsed) the size of P is given by *S*_*j*_(1 − *e*^−*mτ*^)^26^.

Consumption and resource parameters were used as follows. In all simulations 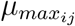 was sampled from a uniform distribution with a lower bound of 0.4 and an upper bound of 1. This equates to a maximum doubling time of 42-104 minutes when resources are non-limiting. In the first set of simulations 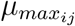 was sampled from a uniform distribution with the same lower and upper bounds as for 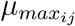. In the second set of simulations *Ks* = *µ*_*max*_^*4*^. In all simulations *Q*_*i*_*j* = 0.1, *S*_*j*_ = 5, *m* = 0.25. Simulations were run with a fixed time step of 0.1 h (dt = 0.1) for 4000 “hours” and initial densities/concentrations of 10 and 5 for all consumers and resources respectively. All models were simulated with the rescomp package (v0.1.0) in R (version 4.1.2).

## Acknowledgements

The authors thank Jan Engelstaedter and Po-Ju Ke thoughtful discussions and comments on an earlier draft of this manuscript.

## Competing Interests

The authors declare no competing financial interests.

## Author contributions

ADL conceived the study and performed simulations. ADL and WBL wrote the manuscript.

## Notes

### Competing Interest Statement

The authors have declared no competing interest.

